# Structure of the fission yeast actomyosin ring during constriction

**DOI:** 10.1101/194902

**Authors:** Matthew T. Swulius, Lam T. Nguyen, Mark S. Ladinsky, Davi R. Ortega, Samya Aich, Mithilesh Mishra, Grant J. Jensen

## Abstract

Cell division in many eukaryotes is driven by a ring containing actin and myosin. While much is known about the main proteins involved, the precise arrangement of actin filaments within the contractile machinery, and how force is transmitted to the membrane remains unclear. Here we use cryosectioning and cryo-focused ion beam milling to gain access to cryo-preserved actomyosin rings in *Schizosaccharomyces pombe* for direct three-dimensional imaging by electron cryotomography. Our results show that straight, overlapping actin filaments, running nearly parallel to each other and to the membrane, form a loose bundle of approximately 150 nm in diameter that “saddles” the inward-bending membrane at the leading edge of the division septum. The filaments do not make direct contact with the membrane. Our analysis of the actin filaments reveals the variability in filament number, nearest-neighbor distances between filaments within the bundle, their distance from the membrane and angular distribution with respect to the membrane.

**Significance Statement:** Most eukaryotic cells divide using a contractile actomyosin ring, but its structure is unknown. Here we use new specimen preparation methods and electron cryotomography to image constricting rings directly in 3D, in a near-native state in the model organism *Schizosaccharomyces pombe*. Our images reveal the arrangement of individual actin filaments within the contracting actomyosin ring.

## Introduction

Cytokinesis, the final step of cell division in eukaryotic cells, is typically driven by a contractile actomyosin ring (AMR) primarily composed of actin (1) and myosin (2). Our understanding of the molecular mechanisms of cytokinesis is most detailed in the rod-shaped unicellular eukaryote *Schizosaccharomyces pombe* (otherwise known as fission yeast), which shares a remarkably conserved set of cytokinesis genes with metazoans (3). In *S. pombe*, the AMR undergoes multiple phases known as assembly, maturation, constriction and disassembly (4), with open questions in each of these four stages. Due to a lack of information about the precise arrangement of filamentous actin (F-actin) within the force-generating network of the AMR, we chose to focus on imaging the AMR during constriction.

In *S. pombe*, glancing sections through plastic-embedded, dividing cells gave the first glimpse of actin filaments running parallel to the division plane at the front of the septum (5). Unfortunately, the study yielded limited examples and lacked three-dimensional (3D) information for a full analysis. In an ambitious pioneering effort, Kamasaki et al., produced 3D reconstructions of entire *S. pombe* AMRs by imaging serial sections through permeabilized cells decorated with Myosin-S1 fragments (6). The amount of F-actin and the size of the rings appeared significantly altered by the procedure used for preserving them (details in discussion), but the continuous bundles that were reconstructed were composed of mixed polarity filaments running circumferentially around the cell.

Here we sought to visualize the precise arrangement of F-actin within the AMR and its interface with the membrane by imaging intact cells in a cryo-preserved state using electron cryotomography (ECT) (7). Because whole *S. pombe* cells are too thick for ECT, which is limited to specimens thinner than a few hundred nanometers, we overcame this obstacle by first rapid-freezing dividing cells and then either cryosectioning them or using the recently-developed method cryo-focused ion beam (FIB) milling to produce thin sections or lamellae suitable for ECT analysis. In this study, ~200-nm wide bundles of straight, overlapping F-actin were seen “saddling” the septum, but no direct contact between filaments and the membrane were observed over 3 μm of total AMR circumference. 3D segmentations of the filaments and membrane allowed for quantitative analysis of the average filament length and number per ring, their persistence length, nearest neighbor distances between filaments as well as between filaments and the membrane. Additionally, the angular distribution of filaments and spatial distribution of filament ends were analyzed. Due to the novelty of the methods and three-dimensional nature of data presented, we urge the reader to first watch Movie S1 for a visual summary of both the methods and main results.

## Results

### ECT of *S. pombe* division sites

To enrich for cells undergoing cytokinesis, we synchronized cells of a temperature sensitive mutant of *S. pombe* (*cdc25*-22 *rlc1*-3GFP) expressing a GFP-tagged regulatory light chain of myosin (Rlc1-3GFP). Transient inactivation of the mitotic inducer phosphatase Cdc25p is a commonly employed approach for synchronization (6, 8, 9), and a detailed characterization of this mutant can be found in the supplementary information (Fig. S1 and Experimental Methods). The use of *cdc25*-22 *rlc1*-3GFP allowed us to monitor the formation of fluorescent cytokinesis nodes and their coalescence into a continuous fluorescent ring near the middle of each cell’s length. Once a majority of the rings had begun to contract (Fig. 1A; Movie S1 at 0:18), cells were vitrified and thinned by either cryosectioning or cryo-FIB milling. Cryosections (top of Fig. 1B, Movie S1 at 0:46) and cryo-FIB milled lamellae (bottom of Fig. 1B; Movie S1 at 1:15) were inspected in a cryo-transmission electron microscope (TEM), and division sites with a visible septum were targeted for tilt-series collection and tomographic reconstruction (Fig. 1B–E; Movie S1 at 1:50).

**Figure 1:**
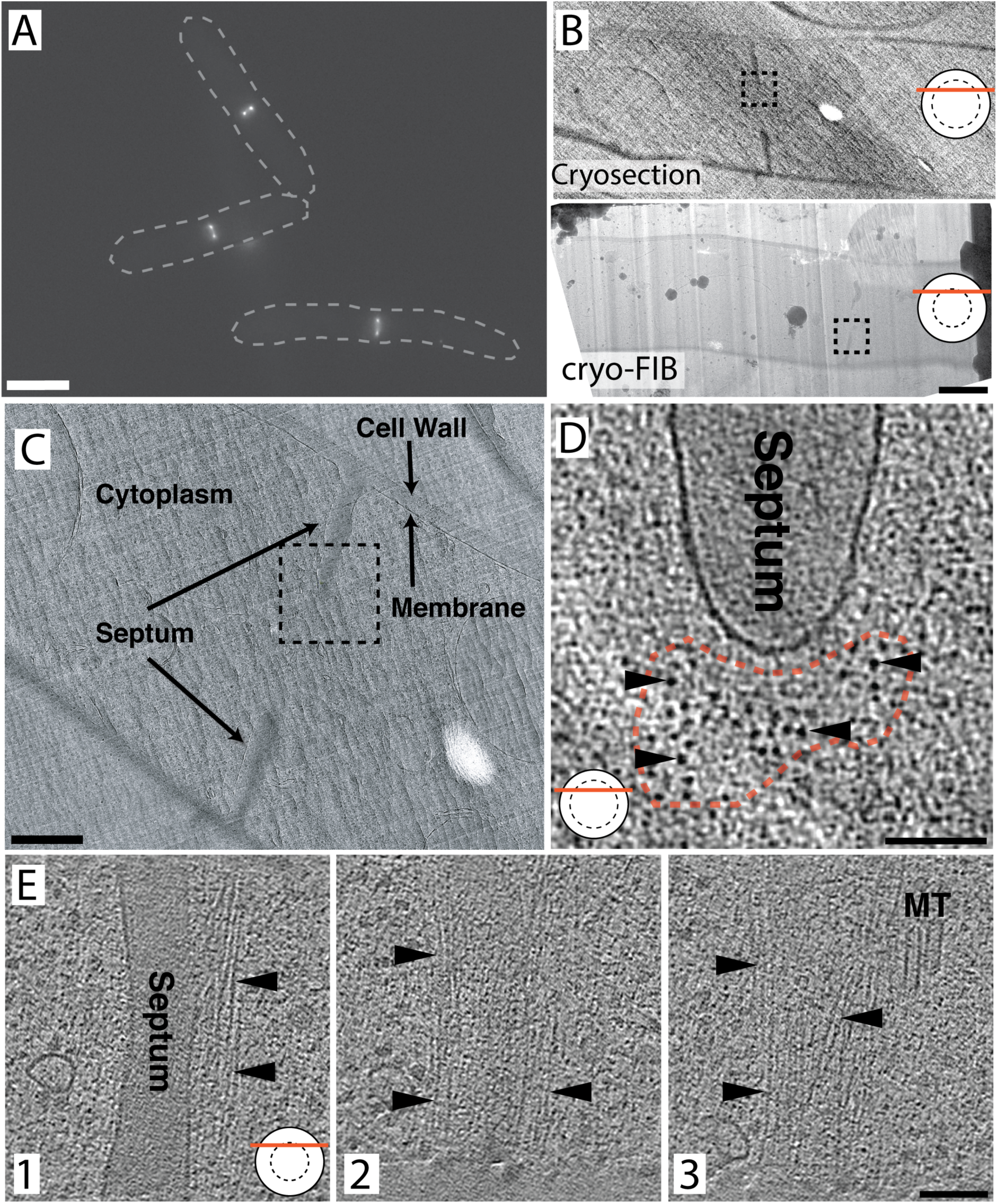
Imaging the AMR by ECT. (A) Fluorescence image of synchronized, contracting AMRs. Dashed lines indicate outlines of whole cells for reference. (B) Projection images of a vitreous cryosection (top) and a cryo-FIB milled lamella (bottom). The boxed region in the top and bottom panel indicate the location of the tomogram in (D) and (E), respectively. (C) Higher mag projection image of the cryosection through a dividing cell in panel B. The boxed region indicates the location of the tomogram in (D). (D) Tomographic slice from a transverse cryosection through the AMR, as indicated by the graphic in the bottom left corner (the solid circle represents the cell in cross-section, the dashed circle represents the ring, and the red line represents the plane of the section through the cell). The red dashed line marks the cross-sectional shape of the actin bundle and arrowheads point to cross-sections through individual actin filaments. (E) Tomographic slices from a FIB-milled lamella cutting through the upper portion of the AMR. Slice 1 shows the leading edge of the septum and slices 2 and 3 are progressively further toward the cytoplasm. Arrowheads point to actin. Two sections of microtubules (MT) are visible. Scale bars represent 5 mm in panel (A), 1 mm in (B), 500 nm in (C) and 100 nm in (D) and (E). Tomographic slices are 20 nm thick.

In total, ~80 tomograms of division sites were generated, and filamentous structures were distinguishable at the leading edge of the division septum in all of them. Sections cut or milled, simply called sections from now on, through the division plane nearer to the central region of the ring produced transverse cross-sections of the contractile ring with putative actin filaments running at small angles with respect to the electron beam. From this perspective, a cluster of densities (or spots) was identifiable near the front edge of the septum (Fig. 1D; Fig. 2A–B; SI Appendix/Fig. S2) in tomographic slices (slices of the 3D reconstruction). Scrolling up and down along the Z-axis of these tomograms revealed that these spots were cross-sections through filaments that typically traversed the thickness of the entire section, though some of them terminate within the section (see Movie S1 at 2:29 and red dots in Movie S2 or Fig. 4B). Note that while small dense spots corresponding to globular proteins were also seen throughout the cytoplasm, they were not continuous across multiple slices. In seven cases, more tangential sections through a region near the top or bottom of the ring were reconstructed in which filaments running perpendicular to the electron beam were visible (Fig. 1E, panels 1–3). As in the transverse sections, filaments in the tangential sections also lined up in a bundle at the leading edge of the division septum (Fig. 1D–E). Note, however, that except for the example in Fig. 1E, all tangential sections captured only portions of the bundle making a thorough analysis from this view implausible.

**Figure 2:**
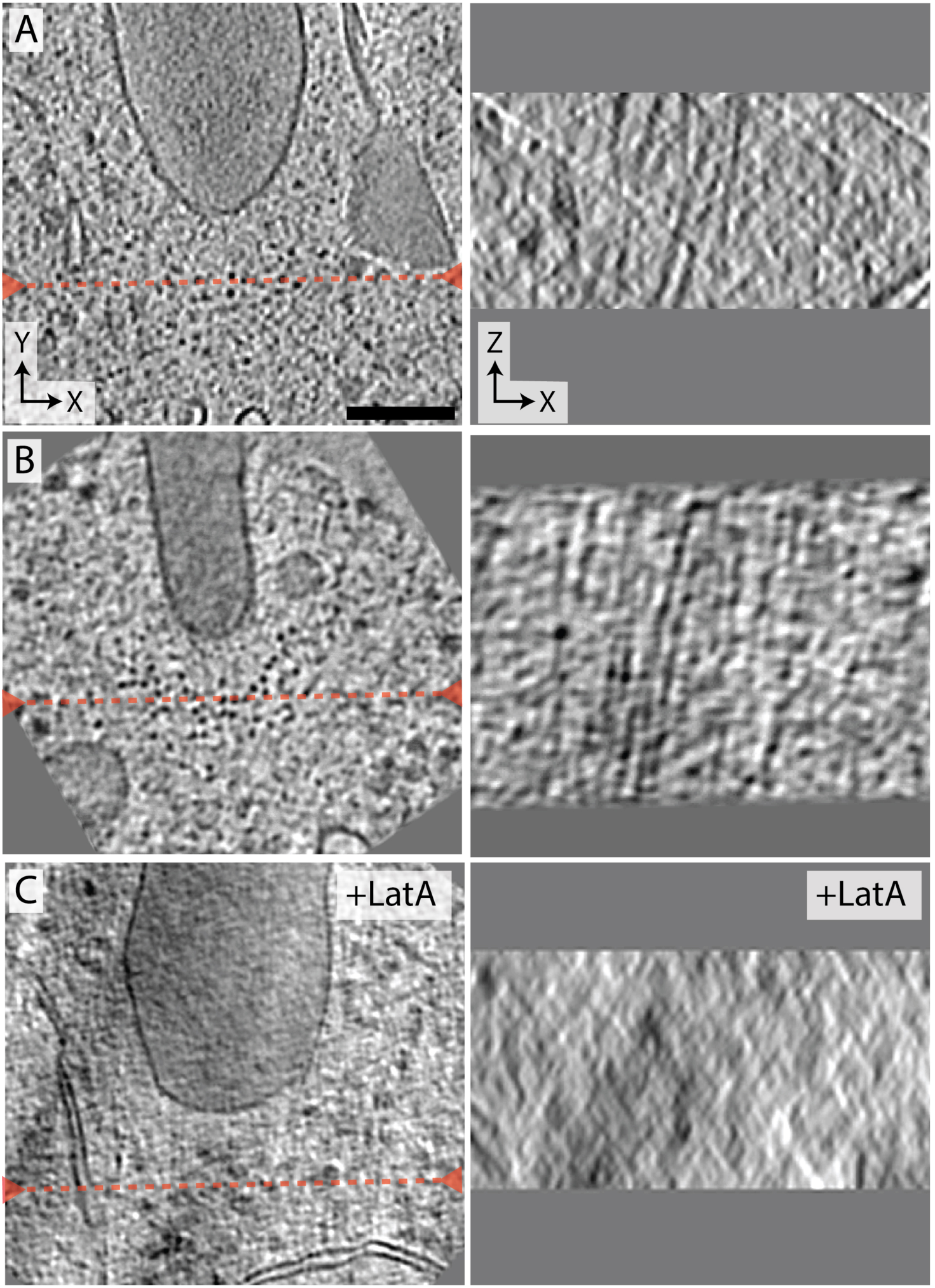
Representative 20 nm-thick slices within transverse sections. The XY slices (left) are perpendicular to the electron beam and the XZ slices (right) are parallel to the beam at the location indicated by the red dashed line. In (A) and (B) where the XY slices show a collection of dots near the leading edge of the septum, the corresponding XZ slices reveal streak patterns indicating the dots are cross-sections of filaments. In (C) where the cells were treated with LatA, the collection of dots and streak pattern are not seen in the XY and XZ slices, respectively. Scale bar represents 100 nm.

### Filaments visualized in the bundle are primarily F-Actin

The molecular identity of each filamentous density within the bundles cannot be easily determined from visual inspection, but from an ultrastructural perspective they appeared to be composed primarily of F-actin. In addition to being filamentous (extending through the depth of the tomogram), they were ~7.5-nm wide, which is consistent with the known size of F-actin. To test this hypothesis, we used ECT on synchronized dividing cells treated with 10 μM Latrunculin A (LatA), which prevents actin polymerization and led to the loss of F-actin in *S. pombe* within 10 min (SI Appendix/Fig S3). In all 6 tomograms collected from LatA-treated cells no filaments were seen at the leading edge of division septa (Fig. 2C & SI Appendix/Fig. S4), further supporting that filamentous densities were F-actin.

We were surprised that no obvious myosin motors could be distinguished by eye in our cryotomograms, and performed correlative light and electron cryomicroscopy (cryoCLEM) on cryosections through dividing *S. pombe* to ensure its presence in the tomograms (Fig. 3; Movie S1 at 3:02). Using the same GFP-tagged regulatory light chain of myosin II (Rlc1-3GFP), all septa seen in the cryosections (five total), by fluorescence cryomicroscopy, correlated with bright GFP puncta (Fig. 3A & D), suggesting that myosin II is present in all of our sections through the AMR. Additionally, tomograms collected on these septa appeared identical to all of our previous tomograms with only F-actin filaments obviously visible (Fig. 3E).

**Figure 3:**
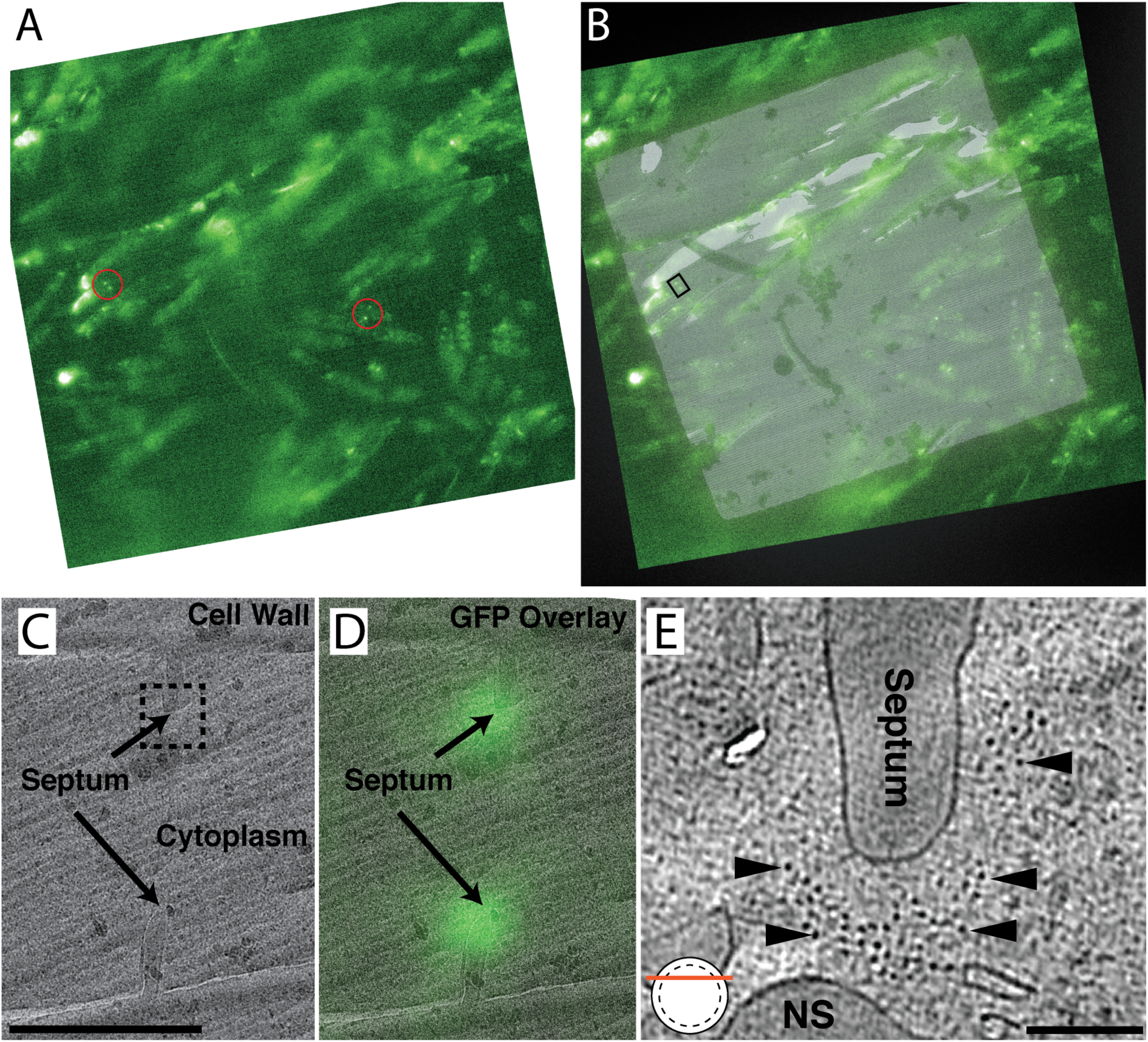
Correlated light and electron microscopy of cryosections of dividing *cdc25-22 rlc1-GFP* cells. (A) A single cryo-fluorescence light microscopy (cryo-FLM) image from a focal stack through one EM grid square that is covered by a vitrified section of a high-pressure frozen pellet of synchronized dividing cells. Red circles highlight two pairs of sharp fluorescent puncta indicative of cross-sections through the myosin-containing contractile ring. (B) Overlay of the fluorescent image and a low-magnification cryoEM image of the cryosection. (C) Higher-magnification image of the region indicated by the small box in panel B, where a division plane is clearly visible. The boxed region indicates the location of the tomogram in panel E. (D) GFP overlay showing the fluorescent puncta are at the leading edge of the septum. (E) Tomographic slice from a transverse cryosection through the AMR, as indicated by the graphic in the bottom left. The red dashed line marks the cross-sectional shape of the actin bundle. Some non-septal (NS) material, which appeared identical to the cell wall in density and texture, is seen in the cytoplasm. Scale bar represents 1 µm in panel C and 100 nm in panel E. The tomographic slice in E is 20 nm thick.

### Analysis of the F-actin Bundle

#### General Characteristics

To analyze AMR filament bundles and their relationship to the membrane, 22 near-to-transverse sections through the AMR (as in Fig. 3E) were segmented as shown in Figure 4. The plasma membranes were segmented by hand, while the central line of each actin filament was segmented computationally using an actin-segmentation algorithm within the Amira software package (10). Using these 3D segmentations, we counted the number of filaments in each tomogram. On average there were 34 (SD = 12) filaments per bundle cross-section with individual numbers ranging between 14 and 60 filaments (Fig. 5), which was largely compatible with quantitative fluorescence microscopy that estimated there would be ~50 filaments per ring cross-section (11). The majority of filaments traversed the entire thickness of the tomographic section, but many filament ends were visible as well (arrowheads in Fig. 4B; Movies S2 and S4). From these 3D segmentations, it was obvious that the cross-sectional boundary of the filament bundle and the distribution of filaments within the bundle varied (Fig. 5). A few tomograms from different sections through the same cell were generated (five pairs marked by asterisks in Fig. 5). While the numbers of filaments were the same in one pair (Fig. 5 O & P), they differed in one pair (Fig. 5 H & I) by a single filament (30 vs. 29), and the other three pairs (Fig. 5 A & B; L & M; U & V) differed more substantially (20 vs. 29; 29 vs. 35; 55 vs. 42). Top-views of the 3D segmentations also revealed variance in the cross-sectional shape of the filament bundle in all five of these cases. Further, filament bundles were pleomorphic and individual filaments did not adhere to rigid lateral constraints like those observed in the sarcomere of muscle (12). In nearly half the cases (10 of 22), the filaments within the bundle cross-section could be divided into “sub-bundles” (SI Appendix/Fig. S6; Movie S3) separated by gaps at least 22 nm wide, which is too far apart to be connected by either the fission yeast actin crosslinker *α*-actinin or fimbrin (we estimated the length of *S. pombe α*-actinin to be 22 nm by combining the length of two actin-binding domains (5 nm each) and two spectrin repeats (6 nm each) estimated from the PDB structure 4D1E). It’s probable that these sub-bundles are actually cross-linked to the AMR in the regions of the ring above or below our ~200-nm thick sections, and it’s also possible that other longer actin-crosslinkers connect these sub-bundles.

**Figure 4:**
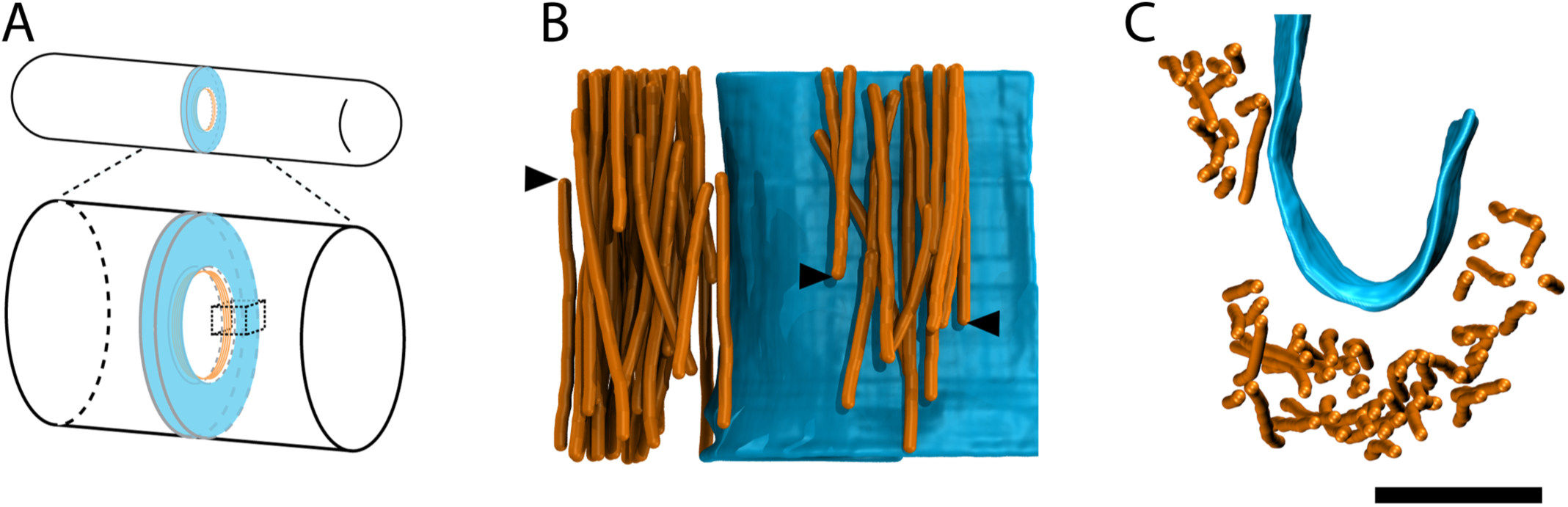
3D segmentations of tomographic reconstructions. (A) Schematic of *S. pombe* division septum (blue) and the AMR (orange) at the leading edge. The dashed cube indicates the portion of the septum represented by the 3D segmentations in panels B and C. (B) Side view of segmented tomographic reconstruction of actin filaments (orange) and the membrane (blue) from a transverse section through the AMR. Arrowheads point to filaments that terminate within the section. (C) Top view of the same segmentation as in panel B. Scale bar represents 100 nm.

**Figure 5:**
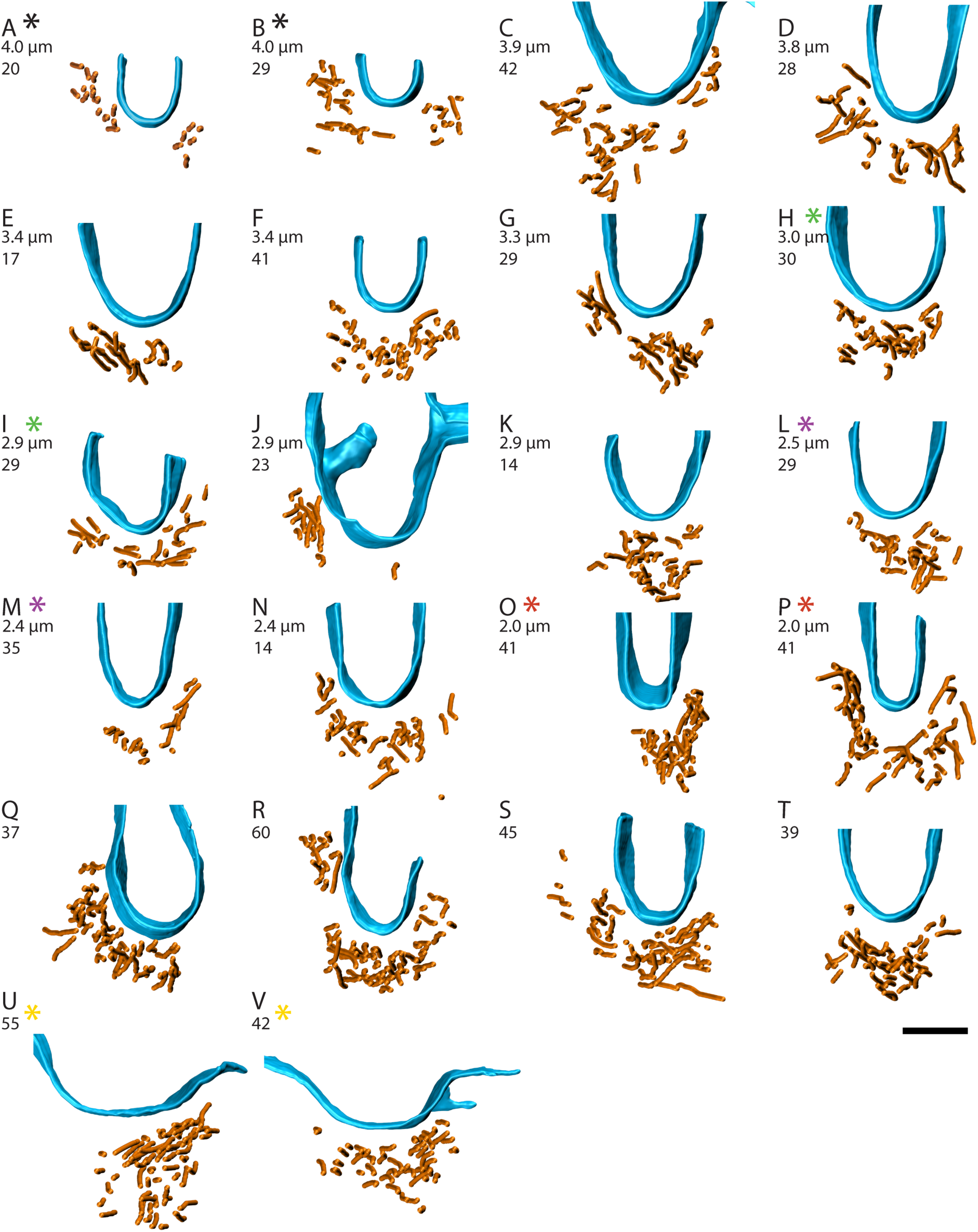
Top views of 3D segmentations. The filaments are in orange and membrane in blue. (A–V) show 22 top views generated from tomographic reconstructions of transverse sections through the AMR, which illustrate the variability in filament number, variability in cross-sectional shape of the AMR, and correlation between membrane curvature and shape of the plasmalemmal face of the AMR. The calculated diameter of the ring is shown just below the panel letters (panels A–P) and the other number (A–V) indicates the number of filaments. Color-coded asterisks represent pairs of reconstructions from the same cell, but from different tomographic sections. Each image corresponds to the same dataset shown in Figure S3. Scale bar represents 100 nm.

#### Estimating Filament Length and Number

Based on the number of filament ends and the thickness of the tomographic sections we estimated that the average length of the filaments was 910 nm (SD = 550 nm) and on average there were 285 filaments in the ring (SD = 180). Note that this calculation depends on the assumption that the protein distribution is uniform around the ring and therefore the filament bundle in each section was representative of the ring. Next, we calculated the ring diameter to determine how these factors depend on the stage of constriction, but as the ring diameter varied we did not see a clear trend in either the average filament length nor the total number of filaments in the ring (Fig. 6A & B). The ring diameter calculation could not be done for six segmented tomograms (Fig. 5 Q–V) due to a lack of corresponding low-magnification images.

**Figure 6:**
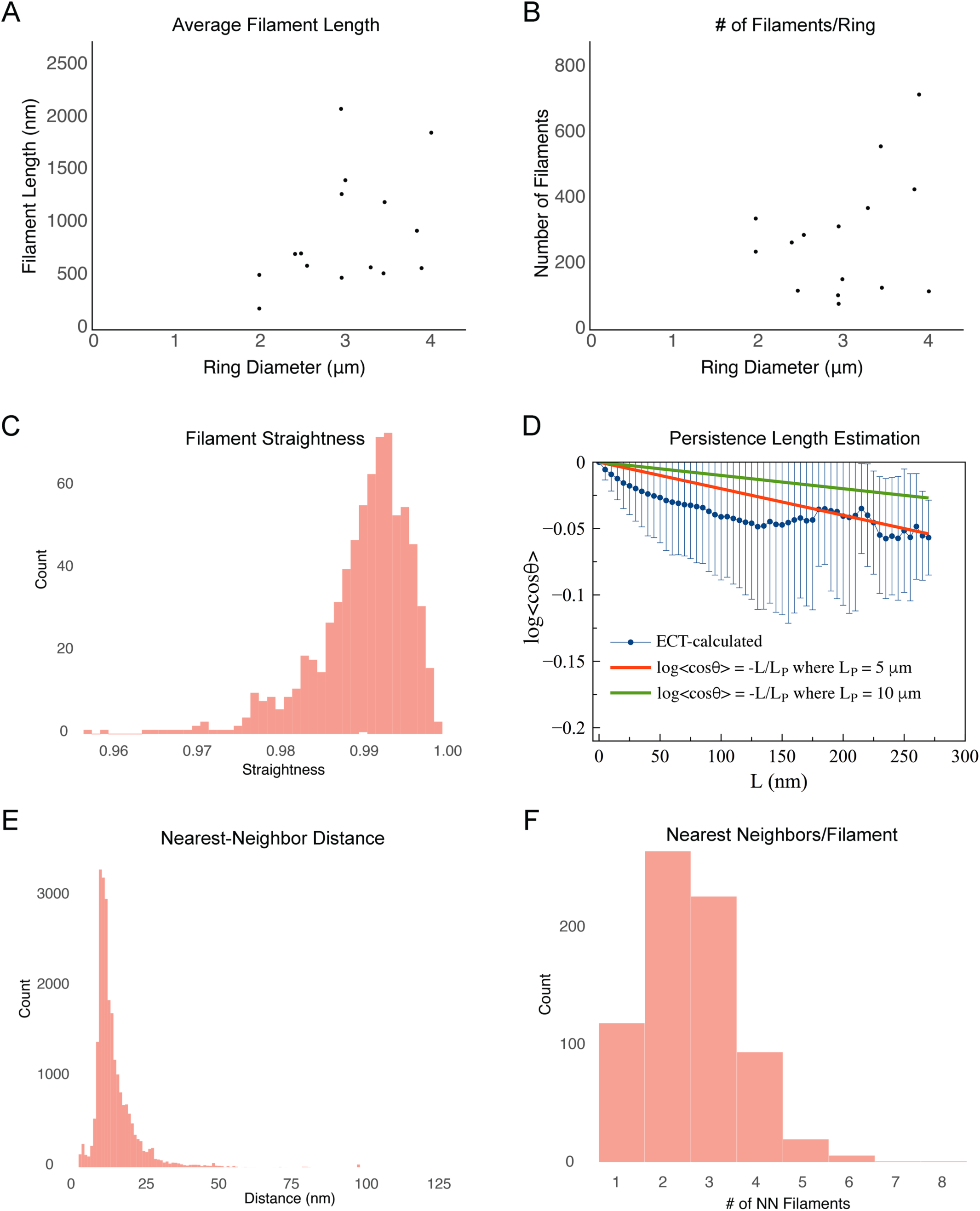
Quantitative analysis of contractile ring filaments. (A) Scatter plot of the estimated filament length as a function of ring diameter. (B) Scatter plot of the estimated number of filaments per ring as a function of ring diameter. (C) Histogram of filament straightness, defined as the ratio of the end-to-end distance to the contour length of the filament (see **Methods** for details). (D) Plot of the persistence length of actin filaments in the tomograms (blue points) compared to the plotted formulas for persistence length equal to 5 μm and 10 μm (the known value for free actin in solution). Error bars indicate standard deviation. (E) Histogram of nearest neighbor distances between the filaments. Each filament was modeled as a chain of beads. Pairs of beads on the same filaments were excluded from the calculation. (F) Histogram of the number of nearest-neighbor partners for each filament in the tomograms. Except for panels A and B, data were combined from all 22 segmented tomograms (740 total filaments). The bin size is 0.1 in (C), and 1 nm in (E).

#### Bundles are Composed of Straight, Nearly Parallel F-actin

From visual inspection of the F-actin bundles in three dimensions (see top and side views in Fig. 5 and SI Appendix/Fig. S5) it appeared that all bundles were composed mainly of straight filaments running nearly parallel to one another. This was quantitatively verified by calculating the straightness of each actin filament as the ratio of its end-to-end distance to its contour length (see schematic in SI Appendix/Fig. S8A), which revealed a narrow distribution of values between 0.97 and 1 (Fig. 6C). Alternatively, a persistence length of ~ 5 μm was calculated from the tomographic data (Fig. 6D), which is half that of free F-actin at ~ 10 μm (13), but more than large enough to appear straight in the ~200-nm thick sections produced for this study.

Next, by representing each filament as a chain of beads, we calculated the distances between all the beads in the bundle except for those pairs that are part of the same filament. The nearest neighbor distance was then determined for every bead along every filament, and the combined data from all 22 tomograms produced a peak that came on sharply at 12 nm and fell off more gradually out to ~50 nm (Fig. 6E), suggesting that most filaments are cross-linked within the bundle. Our analysis was based on center-to-center measurements between filaments, which means that the sharp drop-off below 12 nm translates into a minimum distance of only 4.5 nm between two 7.5 nm-thick filaments. Note that these measurements are in large agreement with an estimate of ~15 nm based on quantitative fluorescence microscopy (11).

Finally, in a perfectly parallel bundle of continuous filaments, the nearest neighbor filament should remain the same throughout a filament’s length. However, when we calculated and plotted the number of distinct nearest neighbor filament-partners for each individual F-actin, we found that individual filaments often had multiple nearest neighbor partners along their length, with a range between 1 and 6, and a peak at 2 (Fig. 6F). This indicates that while the filaments are nearly parallel, there is some degree of intercalation among them.

### The Bundle’s Relationship to the Membrane

The interface between the bundle and the membrane is of particular importance, because force from the ring’s generated tension needs to be transmitted to the ingressing membrane. To better understand this interface, we quantitatively characterized the spatial relationship between the filament bundle and the membrane. First, we calculated the distance between filaments and the membrane by dividing each filament into 20-nm segments and calculating the distance from the midpoint of each segment to the membrane (see schematic in SI Appendix/Fig. S8B). The combined histogram of all 22 tomograms shows a broad peak between 10 and 150 nm, with a mean value of ~60 nm (Fig. 7A), which produces a bundle size (100-200 nm across) that agrees largely with the dimensions observed by Kanbe et al in 1989 and by super-resolution fluorescence (14). Plotting similar histograms for each tomogram reveals the variability in the distribution of filaments with respect to the membrane (SI Appendix/Fig. S7). When the same distance measurements were made between only the F-actin termini and the membrane, a similar distribution of distances was observed (11 nm to 177 nm with a median of 66 nm; Fig. 7B), suggesting that their spatial distributions are determined by the same factors regulating the distance of the filaments as a whole.

**Figure 7:**
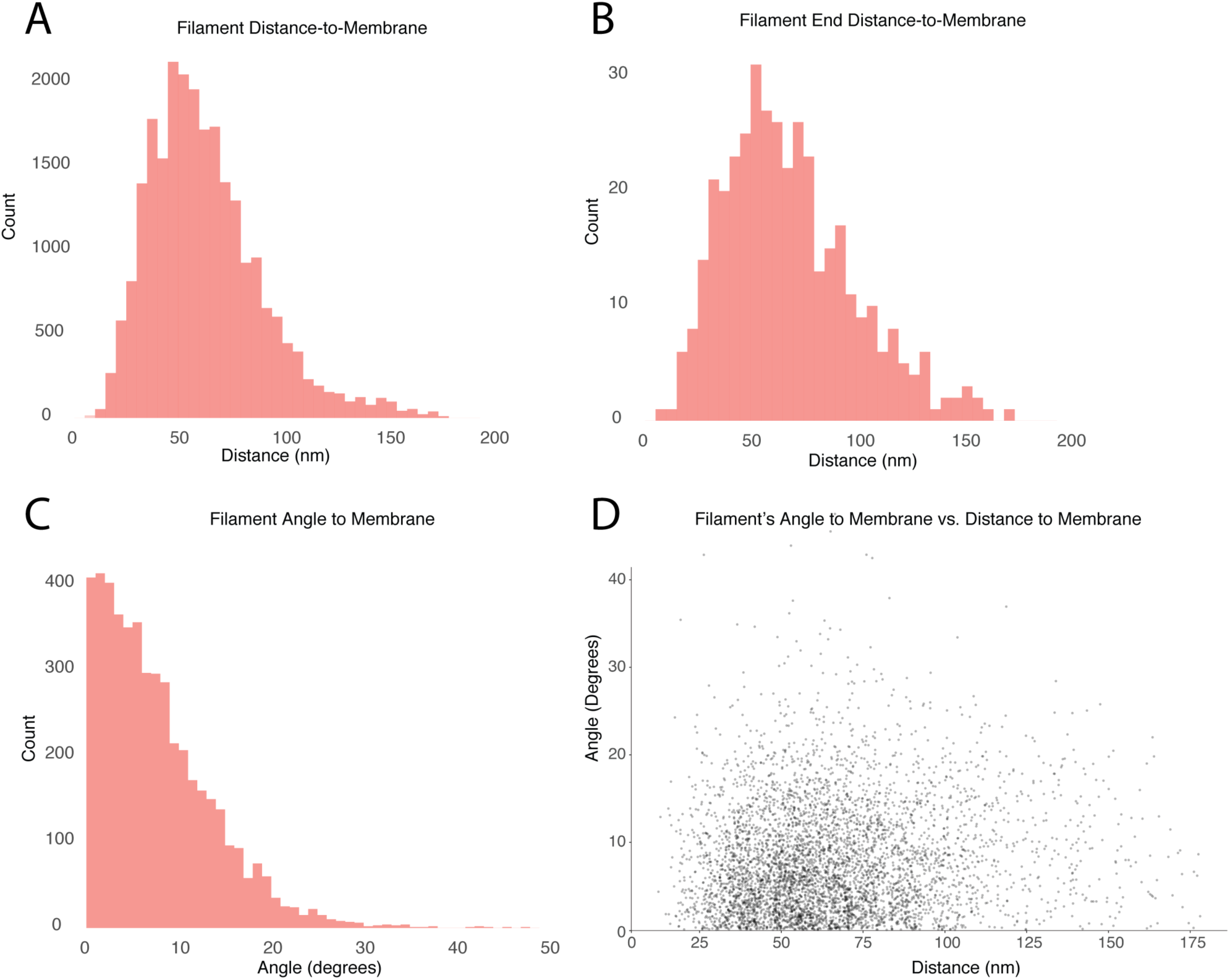
Quantitative analysis of contractile ring filaments with respect to the membrane surface. (A) Histogram of distances measured from the 20-nm segments (N = 5032) of 740 filaments to the membrane. (B) Histogram of the distance measured between filament-ends and the membrane. (C) Histogram of angles between the 20-nm segments and the membrane. (D) Scatterplot of the measured distance from the membrane as a function of the segment’s angle with respect to the membrane surface. The bin size is 1 nm in (A) and (B), and 1 degree in (C).

It appeared by visual inspection of the segmented AMR sections (see top and side views in Fig. 5 and SI Appendix/Fig. S5) that the filaments run nearly parallel to the membrane, and this was verified by calculating the angle between the membrane and each 20-nm long segment of the filaments (see SI Appendix/Fig. S8B). When the combined measurements from all 22 tomograms were plotted as a histogram (Fig. 6C) it showed that nearly all filaments make small angles, with the peak at 2°, and an average of 7.8° across a range from 0° to ~50°, with the majority falling below 20°. It is not clear what governs the angle of the filaments, but to test whether it was a function of the filament’s distance from the membrane, we plotted the angle vs distance and it revealed no correlation (Fig. 7D).

#### The Gap

There is an obvious gap between the filament bundle and the membrane, but it was difficult with our current data to make out any regular structures within this region, regardless of the defocus of the specific tomogram or the thickness of the virtual slice used to view the data. Denoising the tomograms by non-linear anisotropic diffusion (NAD) filtering did not help reveal any regular, interpretable structures within the gap either. Occasionally, a thin density extending from the membrane could be seen, but was either difficult to distinguish from noise or it was rare and unlikely to be responsible for linking the ring to the membrane. The gap did not contain any visible ribosomes, while many were easily seen elsewhere in the cytoplasm, suggesting they are specifically excluded or blocked via steric hindrance from the F-actin or other molecules occupying the gap.

Despite the lack of visible connections to the membrane, the actin bundle’s consistent proximity to the leading edge of the septum, combined with the way it “saddled” the ingressing membrane, implied a physical connection must exist between them (Fig. 5 SI Appendix/Fig. S2; Movie S1 at 4:30). The fact that proteins connecting the actin bundle to the membrane were not visible in the gap suggests an inherent heterogeneity in the architecture of these connections, and that the connections may have been too thin to resolve by our method.

## Discussion

Here we have presented the first pictures of cryo-preserved AMRs, obtained by ECT of cryosectioned or cryo-FIB-milled *S. pombe* cells. General dimensions of the ring, measured by ECT, were in agreement with previous EM data from plastic sections of *S. pombe* (5), and from fluorescence imaging (11, 14). However, the direct imaging of cryo-preserved cells and the three-dimensional nature of electron cryotomography lead to a more detailed understanding of how individual actin filaments are arranged within the ring and their spatial relationship to the membrane.

### Comparison with previous thin-section EM studies

We were surprised by how different the results were from those reported in the previous EM study of spheroplasted, permeabilized, and serially-sectioned cells (6). Apparently, the harsh treatments in the previous study failed to preserve even gross AMR dimensions, since the *pre-constriction* rings were reported to be ~2.5 μm in diameter, which is only about half of the ~4.5 μm one would expect from the diameter of fission yeast. Furthermore, rings from spheroplasted and permeabilized cells were both 10 times wider along the cell’s long axis (based on the thickness and number of serial sections used for reconstruction) and 2–3 times thicker (based on the scale bar shown) (6). The number of filaments in the ring seen here and the previous study also differed significantly: 14–60 filaments per cross-section in vitrified cells versus ~60–180 in the plastic sections (our estimate based on the total number of filaments reported, their average length, and the ring diameter) (6). It is difficult to know, with certainty, why such large discrepancies exist, but given the treatment needed to decorate intracellular F-actin with myosin S1 fragments (cell wall digestion and detergent-permeabilization), it is plausible that the AMRs are no longer under tension because turgor pressure is lost, or that the structure was altered when other binding proteins (especially myosin) were competed away by the myosin S1 fragments. Additionally, membrane permeabilization was conducted in the presence of the actin stabilizing molecule phalloidin, which is known to promote actin polymerization (15) and could account for the increased amount of F-actin in the serial-sectioned rings.

Comparing our ECT results with ultrastructural data from sections of plastic-embedded animal cells (16–18), they share a variety of structural similarities, such as the alignment of filaments parallel to the division plane. In terms of ring dimensions, the ~200-nm thickness of AMRs in animal cells (16, 17) is largely compatible with the thickness measured in our ECT data from fission yeast. As expected, however, the width of the AMR in animal cells is commensurate with its much larger division furrow, which results in AMRs that are many microns in width (16, 17). In the cleavage furrow of sea urchin eggs the ring was described as a uniform band of circumferentially-aligned filaments (16), however, multiple small bundles of actin were observed in HeLa cells (17). These differences might represent natural variation in AMR architecture across cell types or perhaps reflect different preparation methods, but it is possible that the single bundle seen here in yeast by ECT represents some basic unit for cell contractility. To this point, the dimensions of the individual bundles described in HeLa cells (~150 nm in diameter) and their average number of filaments in cross-section (25 filaments) (17) reasonably fit our ECT data of AMRs in *S. pombe* (~100 to 200-nm thickness with 34 filaments on average). Short 13-nm wide filaments, presumed to be myofilaments, were described in HeLa cell AMR bundles, however, and no such myofilament was seen in our tomograms.

While there is a remarkable conservation in the cytokinetic genes of *S. pombe* and animal cells (19), and some of the differences in ring dimensions between fission yeast and animal cells are simply due to differences in cell size, there are also fundamental differences in the way rings assemble and function in *S. pombe* compared to animal cells. For instance, *S. pombe* have a cell wall that provides cell rigidity and shape in the presence of high turgor pressure. There is evidence that cell wall growth in fission yeast actively contributes to constriction force, and is required to generate forces large enough to counter the outward force of turgor pressure (20). Animal cells, in contrast, depend on internal cytoskeletal reinforcement to maintain shape (21), and also have a branched actin cortex linked to the cytoplasmic face of the plasma membrane, unlike yeast. Currently, evidence suggests that AMRs in animal cells are formed from remodeling this cortical actin, which is already present at, and linked to, the membrane (18, 22). Direct 3D imaging by ECT in a range of cell types will be required to compare the properties and intrinsic variability of AMR structures in different systems.

### The structure of myosin II in *S. pombe* actomyosin ring

Our current ECT data was not able to resolve the structure of myosin motors in the AMR, but correlative fluorescence data (Fig. 3) showed that myosin was present in our tomograms, suggesting that neither myosin II isoform in *S. pombe* (Myo2p or My2p) forms the 30-nm thick myofilaments known to exist in muscle sarcomeres (23). This is a simple logical deduction from the fact that such myofilaments were readily distinguishable in transverse cryosections through vitrified muscle in previous studies (24). The structure of purified Myo2p and Myp2p have not been studied by electron microscopy, but myosin II purified from *Acanthamoeba* self-assembles into a range of oligomeric states, with different thicknesses, through tail-to-tail interactions (25, 26). If myosin II in *S.* pombe does not form a thick myofilament, it might adopt one of the other *in vitro* structures observed, such as a simple unipolar or bipolar structure (made of one or two myosin dimers, respectively), or the intermediate-sized 7-nm thick “minifilament”, composed of 16 myosins (25, 26). The 7-nm thickness is consistent with the 7.5-nm thick filaments seen in the AMR, meaning a subset of the filaments seen could be myosin minifilaments. We currently favor the interpretation that all the filaments we saw were F-actin, however, because no clusters of myosin heads, like those observed in vitro (25, 26), were seen. It is possible, however, that such details were obscured by the relatively close packing of filaments in the bundle.

### Crosslinking in the bundle

Even though we could not confidently identify the physical connections between filaments in the bundle, the relatively tight packing of the filaments suggests that actin crosslinkers were present. Nearest neighbor distance analysis resulted in a peak at 12 nm, corresponding to a gap of 4.5 nm between the 7.5 nm-thick filaments. This distance is much shorter than both fimbrin and *α*-actinin, two known actin crosslinkers that are present in the ring (27), but it’s possible that some unknown crosslinkers of ~ 4.5 nm length may also be present in the ring. It’s also possible that the tension between filaments, generated by myosin, pulled the filaments as close as they could be to one another, given the flexibility of longer actin cross-linkers like *α*-actinin and fimbrin.

### Connecting the AMR to the membrane

A major question that still remains is how the AMR transfers contractile force to the membrane during constriction? Our tomograms revealed a clear gap between the bundle and the membrane with no obvious connecting densities between the actin filaments and the membrane. Additionally, over 3 μm of cumulative AMR length, no actin filaments made direct contact with the membrane. That being said, it seems highly likely that there is some physical connection between the bundle and the membrane or it would not stay localized to the tip of the ingressing septum, nor would its cross-sectional shape saddle the curved edge of the septum with such consistency, as in our observations.

The most obvious potential contact points between the ring and membrane are the cytokinetic nodes that function to recruit and move the major components of the AMR to the division plane during assembly (14, 28–30). These nodes are membrane bound and they contain Myo2p, which interacts with F-actin directly. They have been shown to persist within the ring during constriction by super-resolution live cell fluorescence (14). There are ~140 nodes per cell, so even assuming the largest diameter ring in our experiments (4.0 μm) there would still be one node every 90 nm of AMR on average. That means we would have captured 1-3 nodes per tomogram (100-300 nm thick). If nodes concentrate as the cell constricts then we would have captured even more in our tomograms.

In 2016, Laplante et al. proposed a model of the average cytokinetic node based on both the stoichiometry and spatial arrangement of proteins within the node measured by super-resolution live-cell fluorescence (14). Their model node included four Mid1p dimers, four Cdc12p dimers, 8 Rng2 dimers, 8 Cdc15p dimers and eight Myo2p dimers connected to the node “core” by their coiled-coil tails. The node core (excluding the thin, flexible Myo2p molecules radiating from it) would be ~5.2 megadaltons with an estimated size of ~50 nm x 50 nm x 50 nm, which is more massive but significantly larger than the 80S ribosomes (3.2 megadaltons and ~30 nm x 20 nm x 20 nm) that are clearly visible in the cytoplasmic regions of the tomograms. Presuming the dimensions of the cytokinetic model node is accurate, the 80S ribosome is ~ 5-fold denser than the cytokinetic node (0.27 kDa/nm^3^ vs. 0.05 kDa/nm^3^, respectively), which could account for why such a large complex was undetectable.

The average distance between the filaments and the membrane in our cryotomograms (~60 nm), is consistent with the dimensions of the model node presented by Laplante et al. (14), and their presence could account for the gap we see between the F-actin bundle and the membrane. It could be that during constriction the nodes became so densely packed that individual node assemblies could not be distinguished in the cryotomograms, but the gap between the filament bundle and the membrane appears more similar to the cytoplasmic background than a region packed tightly with protein. Also, statistical analysis of 20×20×100 voxel subvolumes from within the gap showed only a 10% increase in the mean pixel value compared to cytoplasm in the same tomogram. Another possibility is that nodes are not so well-ordered, nor stable during constriction and they become more “fluid”, repeatedly breaking apart and reassembling under competing forces from Myo2p motors connected to the node and the AMR. This kind of instability would make for a very heterogeneous and disordered protein distribution in the gap, which would be consistent with the density seen in the tomograms.

Another possibility is that the distribution of protein mass within the nodes is significantly different than the model presented by Laplante et al., and the bulk of the protein is, for instance, distributed more evenly along the surface of the membrane, making it difficult to detect by ECT. In this case, it would be feasible to think the filament bundle is tethered to the membrane along its length by protein linkers that are simply too thin to be resolved in a cryotomogram, which typically attain 3-4 nm resolution (7). This linker would need to be at least ~27 nm long to connect the bundles in the AMR to the membrane directly, because that is the average nearest distance between the membrane and the closest filament in each of the 22 tomograms analyzed (SI Appendix/Fig. S7). Again, the most viable candidate that meets these requirements is Myo2p, which was suggested by fluorescence microscopy studies of ring assembly to exist as a unipolar myosin (14, 30), and aptly, its coiled-coil tail would have been difficult to resolve in our tomograms. Interestingly, Myo2p is essential for cytokinesis in *S. pombe* (31, 32), even in mutant strains where the motor activity of Myo2p has been greatly reduced (33), suggesting that it plays a critical role in some other function, such as linking the AMR to the membrane.

The node protein formin Cdc12p nucleates and elongates unbranched F-actin from its plus-end (34). Much of the actin in the ring is Cdc12p-derived (35, 36), so it has been proposed that actin filaments might remain engaged with node-bound Cdc12p at their plus ends, thus connecting filaments to the membrane (14, 30). This arrangement is logically satisfying, because myosin II walks preferentially toward the actin plus-end, meaning that any myosin II motor bound to the filament would be putting tension on the membrane. Our tomographic data does not support a model where most filament ends are engaged with node-bound Cdc12, because given that there are 2-4 Cdc12 dimers per node (35, 37), 20-60% of filament plus-ends (half the total ends observed) would not be bound to Cdc12p. Super-resolution fluorescence localization data showed that Cdc12p exists in nodes ~45 nm away from the membrane (14), but the broad peak of filament-end to membrane distances observed by ECT (Fig. 7B) shows that the filament ends to the left of the peak do not cluster at 45 nm. If the majority of filament ends were bound to Cdc12p, one would expect to see a peak around this distance. For now, exactly how the ring is bound to the membrane remains unclear.

## Experimental Methods

### Cell Synchronization, fluorescence imaging, and latrunculin A treatment

To maximize the probability of cryosectioning/FIB milling though actively constricting division septa, a temperature sensitive mutant of *S. pombe* (*cdc25*-22 *rlc1*-3GFP) carrying a GFP-tagged regulatory light chain of myosin II was used. First, colonies grown on YES-agar plates at the permissive temperature (22° C) were harvested with a sterile loop and suspended in liquid YES media to an OD_600_~ 0.1 and further grown at the permissive temperature with 180 rpm shaking. The cells were grown for ~24 hours and kept between OD_600_ = 0.1–0.5. The synchronization process was initiated at OD_600_ = 0.25 with transfer to the restrictive temperature (36° C) for 3.5–4 hours with shaking. Once the OD doubled to 0.5 cells were inspected by light microscopy to ensure that most cells had more than doubled in length due to stalled entry into mitosis. At this point, the cells were shifted to the permissive temperature (22° C) and allowed to enter mitosis.

To prepare samples for ECT, around 45 min after the cells entered mitosis they were screened by epifluoresence microscopy on a Nikon Eclipse 90i microscope using a 100x oil objective (NA = 1.4) and a Photometrics CoolSnap HQ2 CCD for the early formation of rings at the mid-cell. Cells were monitored every 5–10 minutes until ~90% of cells contained intact rings that had begun to contract. Finally, ~5 ml of synchronized culture was either pelleted at 4° C and placed on ice for high pressure freezing (HPF) or un-pelleted cells were directly plunge-frozen on EM grids.

To disrupt F-actin within dividing cells the same synchronization process was used, but during ring constriction latrunculin A (LatA, molecular probes #L12370) was added to the culture to a final concentration of 10 μM. Within 10 min the continuous fluorescent ring became fragmented into puncta along the membrane around the mid-cell. Cells were then pelleted for HPF as described above.

### Characterization of cytokinesis in the temperature-sensitive Cdc25 mutant

To ensure phenotypically normal cytokinesis in the temperature-sensitive *cdc25-22 S. pombe* mutant, we synchronized cells expressing a GFP-tagged regulatory light chain of myosin (Rlc1-3GFP) and an mCherry-tagged spindle pole protein (mCherry-Pcp1p) at the G2-M boundary. Transient inactivation of mitotic inducer phosphatase Cdc25p is a commonly employed approach for synchronization (6, 8, 9). After release to permissive temperature, elongated *cdc25*-22 cells undergo phenotypically normal cytokinesis (SI Appendix/Fig. S1). Rlc1 formed a uniform ring like that of the wild-type cells (SI Appendix/Fig. S1A). Ring assembly and ring constriction started at the onset and the end of spindle pole body (SPB) separation (SI Appendix/Fig. S1B) respectively, similar to what occurs in wild-type cells (38). Though we did observe a modest delay in AMR formation (SI Appendix/Fig. S1C), the time taken for completion of cytokinesis was comparable to those of wild-type cells (SI Appendix/Fig. S1D). Importantly, our method of synchronization had no significant impact on the velocity of ring constriction (SI Appendix/Fig. S1E & F). Thus, synchronization by reversible heat inactivation of mitotic inducer Cdc25p does not significantly alter the dynamics of AMRs.

To characterize cytokinesis in *cdc25*-22 *rlc1*-3GFP, mid-log phase cells were spotted on a 2% Agar pad supplemented with YES media and observed under a custom built spinning disk confocal with an inverted Olympus IX-83,100X/1.4 plan-apo objective, a deep cooled Hamamatsu ORCA II–ER CCD camera and Yokogawa CSU:X1 spinning disk (Perkin-Elmer). A stack of 18–20 Z slices of 0.3 mm Z-step-size was collected every 2 min for an hour at 25° C using the Velocity software (Perkin-Elmer). Images were then rotated and cropped using the imageJ software to align cells and 3D reconstruction was done using the Velocity software.

### Cryosectioning and Cryo-FIB milling

For cryosectioning, synchronized and LatA-treated *S. pombe cdc25*-22 *rlc1*-3GFP cells (OD_600_ = 0.5) were harvested (7197 ×g, 10 min), and the pellet was mixed with 40% dextran (w/v) in YES media. The samples were transferred to brass planchettes and rapidly frozen in a HPM010 high-pressure freezing machine (Bal-Tec, Leica). Note that, the step of first transferring cells to the location of the high-pressure freezer, then mixing cells with dextran, and finally loading a high pressure planchette took ~10–20 min. During this period cells were kept on ice to slow down constriction and after cryopreservation was completed, unused cells on ice were imaged again to ensure that continuous fluorescent rings were still visible. (We found that precooling cells for 20 min on ice did not affect the ring’s constriction rate measured by fluorescence microscopy and the characteristics of the actin bundle revealed by ECT). Cryosectioning of the vitrified samples was done as previously described (39, 40). Semi-thick (150–200 nm) cryosections were cut at −145°C or −160°C with a 25° Cryodiamond knife (Diatome, Biel, Switzerland), transferred to grids (continuous carbon-coated 200-mesh copper) and stored in liquid nitrogen.

For cryo-FIB milling, 4 μl of synchronized *cdc25*-22 *rlc1*-3GFP cells (OD_600_ = 0.5) were applied to the carbon surface of a freshly glow-discharged copper Quantifoil R2/2 EM grid. Grids were then blotted manually, from the back, within the humidifying chamber (95%) of an FEI Mark IV Vitrobot by setting the blot number to zero and using a large pair of forceps to insert a piece of Whatman #1 filter paper through the open side port.

Plunge-frozen grids were then mounted in custom-modified Polara cartridges with channels milled through the bottom, which allowed samples to be milled at a low angle of incidence (~10–12°) with respect to the carbon surface. These modified cartridges were transferred into an FEI Versa 3D equipped with a Quorum PP3010T Cryo-FIB/SEM preparation system. Samples were sputter coated with 20 nm of platinum prior to milling to minimize curtaining and to protect the front edge of the sample during milling. Vitrified cells lying approximately perpendicular to the FIB beam were located and lamellae (~12 μm wide and ~2 μm thick) were rough-milled at the mid-cell with beam settings of 30 keV and 0.300 nA. A polishing mill at a reduced current of 30 pA was then performed to bring the final thickness to 150–400 nm. Samples were then removed from the scope while maintaining a temperature below −160°C and stored in liquid nitrogen.

### Cryo-fluorescence light microscopy (Cryo-FLM)

Frozen grids with attached cryosections were loaded into Polara EM cartridges, transferred into a cryo-FLM stage (FEI Cryostage decribed in Nickell et al., 2006, modified to hold Polara EM cartridges as described in (41)) and imaged on a Nikon Eclipse Ti inverted microscope using a 60× extra-long-working-distance air objective (Nikon CFI S Plan Fluor ELWD 60× NA 0.7 WD 2.62-1.8 mm) and an Andor CCD. Focal stacks of each grid square covered by cryosectioned material were collected using the GFP filter set. Pairs of fluorescent puncta (corresponding to the two sides of the septum) were located within the cryosections and targeted for ECT after transfer to an FEI Tecnai F30 Polara TEM. Fluorescence images were scaled to the electron micrographs, and then overlaid and registered using the whole-cell background fluorescence from the cryosectioned cells.

### Electron cryotomography (ECT) and image processing

Tilt-series of division septa in both cryosections and cryo-FIB milled lamellae were collected on an FEI Tecnai F30 Polara TEM operating at 300 keV every one degree from −60 to +60° with a total dose of 140– 180 electrons/Å^2^. 4k x 4x images were collected with a Gatan K2 Summit direct electron detector in counting mode. Tilt-series were binned to pixel sizes of 1.2 and 0.9 nm/px depending on their original magnification, and then reconstructed using the patch tracking method included in the IMOD image processing suite (42).

Actin centerlines were computationally extracted using a template matching/tracing algorithm implemented within the Amira image segmentation software (43). Before filament extraction, tomograms were denoised using Nonlinear Anisotropic Diffusion filtering in IMOD. To segment the actin filaments, a 3.5 nm radius surface was generated around the filament centerline to match the actual size of actin filaments. Division septum membranes in each tomogram were hand segmented in Amira. Histograms of nearest distance measurements between segmented filament surfaces and the septum membrane were generated by surface-to-surface measurements within Amira. Nearest neighbor distances between filaments were calculated from the coordinates of the extracted filament centerlines using a custom script.

### Actin Straightness

For each filament we calculated its contour length *L*_*contour*_ (as traced by the actin-segmentation software described above) and the length of a straight line connecting the two ends *L*_*end-to-end*_. Filament straightness was then defined as the ratio *L*_*end-to-end*_/*L*_*contour*_.

### Calculation of persistence length

Theoretically, the tangent correlation 〈*cosθ*〉, where *θ* is the angle between the tangent vector at position 0 and the tangent vector at a distance *L* along the filament, is defined as

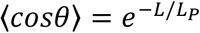

where *L*_*P*_ is the filament’s persistence length. This formula can be expressed as

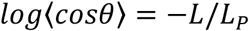

To calculate the persistence length of the segmented filaments, each filament was modeled as a chain of beads with adjacent beads separated by 5 nm. Tangent correlation 〈*cosθ*〉 was then calculated along the beads and *log*〈*cosθ*〉 was plotted vs *L* to derive the persistence length *L*_*P*_. See SI Appendix/Fig. S8A.

### Calculation of ring diameter

The average diameter of our synchronized cells (4.5 μm, SD = 0.18 μm, N = 30) was measured from DIC images collected on a Nikon Eclipse 90i microscope using a 100x oil objective (NA = 1.4) and a Photometrics CoolSnap HQ2 CCD with a detector pixel size of 6.5 μm. For simplicity, the cell diameter was assumed to be 4.5 μm for all cells imaged by ECT. By measuring the width of cell section *A* = 2*a* (SI Appendix/Fig. S9) we could calculate the distance *c* from the center to the section using the Pythagorean theorem as following

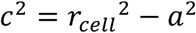

where *r*_*cell*_ was the radius of the cell.

Next, by measuring the distance *B* = 2*b* between the two septal tips in the section we could calculate the ring radius using the Pythagorean theorem:

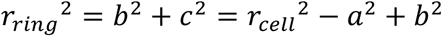

### Average filament length and number of filaments per ring

To calculate the average filament length we assume each tomogram was representative of the ring such that the filaments were uniformly distributed around the ring. As a result, the ratio of filament length to the number of ends was constant and equaled *L*_*a*_/2 for each filament of average length *L*_*a*_ had two ends. By calculating the total filament length *L*_*sum*_ and total number of ends *N*_*e*_ in the tomogram we derived the average length as:

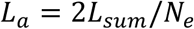

To calculate the total number of filaments in the ring *N*_*fil*_ we calculated the density of filament ends in the tomogram as *N*_*e*_/*T* where *T* was the tomogram thickness. As this density was assumed to be uniform, it was theoretically equal to the ratio of the total number of ends in the ring 2*N*_*fil*_ to the ring length *πD*_*ring*_, or

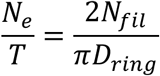

The total number of filaments in the ring was then calculated as:

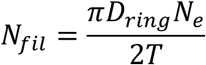

## Acknowledgments

The authors thank Catherine Oikonomou for revising the manuscript for clarity and Frances Allen for initial training and guidance in cryo-FIB milling.

M.M. is an Intermediate Fellow of the Wellcome Trust-DBT India Alliance (IA/I/14/1/501317). M.M. acknowledges the India Alliance and the DAE/TIFR for funds.

## References

1. Schroeder TE (1973) Actin in dividing cells: contractile ring filaments bind heavy meromyosin. Proc Natl Acad Sci USA 70(6):1688–1692.

2. Mabuchi I, Okuno M (1977) The effect of myosin antibody on the division of starfish blastomeres. J Cell Biol 74(1):251–263.

3. Balasubramanian MK, Bi E, Glotzer M (2004) Comparative analysis of cytokinesis in budding yeast, fission yeast and animal cells. Curr Biol 14(18):R806–18.

4. Cheffings TH, Burroughs NJ, Balasubramanian MK (2016) Actomyosin Ring Formation and Tension Generation in Eukaryotic Cytokinesis. Curr Biol 26(15):R719–R737.

5. Kanbe T, Kobayashi I, Tanaka K (1989) Dynamics of cytoplasmic organelles in the cell cycle of the fission yeast Schizosaccharomyces pombe: three-dimensional reconstruction from serial sections. J Cell Sci 94 (Pt 4):647–656.

6. Kamasaki T, Osumi M, Mabuchi I (2007) Three-dimensional arrangement of F-actin in the contractile ring of fission yeast. J Cell Biol 178(5):765–771.

7. Gan L, Jensen GJ (2012) Electron tomography of cells. Q Rev Biophys 45(1):27–56.

8. Nurse P (1975) Genetic control of cell size at cell division in yeast. Nature 256(5518):547–551.

9. Russell P, Nurse P (1986) cdc25+ functions as an inducer in the mitotic control of fission yeast. Cell 45(1):145–153.

10. Rigort A, et al. (2012) Automated segmentation of electron tomograms for a quantitative description of actin filament networks. J Struct Biol 177(1):135–144.

11. Courtemanche N, Pollard TD, Chen Q (2016) Avoiding artefacts when counting polymerized actin in live cells with LifeAct fused to fluorescent proteins. Nat Cell Biol 18(6):676–683.

12. Burgoyne T, Muhamad F, Luther PK (2008) Visualization of cardiac muscle thin filaments and measurement of their lengths by electron tomography. Cardiovasc Res 77(4):707–712.

13. Isambert H, et al. (1995) Flexibility of actin filaments derived from thermal fluctuations. Effect of bound nucleotide, phalloidin, and muscle regulatory proteins. J Biol Chem 270(19):11437–11444.

14. Laplante C, Huang F, Tebbs IR, Bewersdorf J, Pollard TD (2016) Molecular organization of cytokinesis nodes and contractile rings by super-resolution fluorescence microscopy of live fission yeast. Proceedings of the National Academy of Sciences 113(40):E5876–E5885s.

15. Cooper JA (1987) Effects of cytochalasin and phalloidin on actin. J Cell Biol 105(4):1473–1478.

16. Schroeder TE (1972) The contractile ring. II. Determining its brief existence, volumetric changes, and vital role in cleaving Arbacia eggs. J Cell Biol 53(2):419–434.

17. Maupin P, Pollard TD (1986) Arrangement of actin filaments and myosin-like filaments in the contractile ring and of actin-like filaments in the mitotic spindle of dividing HeLa cells. J Ultrastruct Mol Struct Res 94(1):92–103.

18. Mabuchi I, Tsukita S, Tsukita S, Sawai T (1988) Cleavage furrow isolated from newt eggs: contraction, organization of the actin filaments, and protein components of the furrow. Proc Natl Acad Sci USA 85(16):5966–5970.

19. Pollard TD, Wu J-Q (2010) Understanding cytokinesis: lessons from fission yeast. Nat Rev Mol Cell Biol 11(2):149–155.

20. Proctor SA, Minc N, Boudaoud A, Chang F (2012) Contributions of turgor pressure, the contractile ring, and septum assembly to forces in cytokinesis in fission yeast. Curr Biol 22(17):1601–1608.

21. Pollard TD, Cooper JA (2009) Actin, a Central Player in Cell Shape and Movement. Science 326(5957):1208–1212.

22. Fishkind DJ, Wang YL (1993) Orientation and three-dimensional organization of actin filaments in dividing cultured cells. J Cell Biol 123(4):837–848.

23. Al-Khayat HA (2013) Three-dimensional structure of the human myosin thick filament: clinical implications. Glob Cardiol Sci Pract 2013(3):280–302.

24. Trus BL, et al. (1989) Interactions between actin and myosin filaments in skeletal muscle visualized in frozen-hydrated thin sections. Biophys J 55(4):713–724.

25. Pollard TD (1982) Structure and polymerization of Acanthamoeba myosin-II filaments. J Cell Biol 95(3):816–825.

26. Sinard JH, Stafford WF, Pollard TD (1989) The mechanism of assembly of Acanthamoeba myosin-II minifilaments: minifilaments assemble by three successive dimerization steps. J Cell Biol 109(4 Pt 1):1537–1547.

27. Wu JQ, Bähler J, Pringle JR (2001) Roles of a Fimbrin and an-Actinin-like Protein in Fission Yeast Cell Polarization and Cytokinesis. Mol Biol Cell 12(4):1061–1077.

28. Wu J-Q, et al. (2006) Assembly of the cytokinetic contractile ring from a broad band of nodes in fission yeast. J Cell Biol 174(3):391–402.

29. Vavylonis D, Wu J-Q, Hao S, O'Shaughnessy B, Pollard TD (2008) Assembly mechanism of the contractile ring for cytokinesis by fission yeast. Science 319(5859):97–100.

30. Laporte D, Coffman VC, Lee I-J, Wu J-Q (2011) Assembly and architecture of precursor nodes during fission yeast cytokinesis. J Cell Biol 192(6):1005–1021.

31. Kitayama C, Sugimoto A, Yamamoto M (1997) Type II myosin heavy chain encoded by the myo2 gene composes the contractile ring during cytokinesis in Schizosaccharomyces pombe. J Cell Biol 137(6):1309–1319.

32. May KM, Watts FZ, Jones N, Hyams JS (1997) Type II myosin involved in cytokinesis in the fission yeast, Schizosaccharomyces pombe. Cell Motil Cytoskeleton 38(4):385–396.

33. Laplante C, et al. (2015) Three myosins contribute uniquely to the assembly and constriction of the fission yeast cytokinetic contractile ring. Curr Biol 25(15):1955–1965.

34. Kovar DR, Kuhn JR, Tichy AL, Pollard TD (2003) The fission yeast cytokinesis formin Cdc12p is a barbed end actin filament capping protein gated by profilin. J Cell Biol 161(5):875–887.

35. Coffman VC, Nile AH, Lee I-J, Liu H, Wu J-Q (2009) Roles of formin nodes and myosin motor activity in Mid1p-dependent contractile-ring assembly during fission yeast cytokinesis. Mol Biol Cell 20(24):5195–5210.

36. Coffman VC, Sees JA, Kovar DR, Wu J-Q (2013) The formins Cdc12 and For3 cooperate during contractile ring assembly in cytokinesis. J Cell Biol 203(1):101–114.

37. Wu J-Q, Pollard TD (2005) Counting cytokinesis proteins globally and locally in fission yeast. Science 310(5746):310–314.

38. Wu J-Q, Kuhn JR, Kovar DR, Pollard TD (2003) Spatial and temporal pathway for assembly and constriction of the contractile ring in fission yeast cytokinesis. Dev Cell 5(5):723–734.

39. Ladinsky MS, Pierson JM, McIntosh JR (2006) Vitreous cryo-sectioning of cells facilitated by a micromanipulator. J Microsc 224(Pt 2):129–134.

40. Ladinsky MS (2010) Micromanipulator-assisted vitreous cryosectioning and sample preparation by high-pressure freezing. Meth Enzymol 481:165–194.

41. Briegel A, et al. (2010) Correlated light and electron cryo-microscopy. Meth Enzymol 481:317–341.

42. Kremer JR, Mastronarde DN, McIntosh JR (1996) Computer visualization of three-dimensional image data using IMOD. J Struct Biol 116(1):71–76.

43. Rigort A, et al. (2012) Automated segmentation of electron tomograms for a quantitative description of actin filament networks. J Struct Biol 177(1):135–144.

